# Analysis of oogenesis defects in *Dyro* mutant of *Drosophila melanogaster*

**DOI:** 10.1101/2024.08.07.607114

**Authors:** Takamoto Shima, Yuuki Kawabata, Yoshimasa Yagi

**Author notes:** These authors contribute equally (Takamoto Shima, Yuuki Kawabata).

## Abstract

*Dyro* mutant is female sterile, and oogenesis is aborted during stage8-9 of oogenesis. We investigated detail of the oogenesis defect of *Dyro* mutant. At first, we confirmed loss of Dyro caused female sterility by genetic rescue experiment. Then, we performed genetic mosaic analysis and found Dyro expression in germ cell is important for oogenesis. In *Dyro* mutant, inhibition of programmed cell death suppressed cell death of germ cells during oogenesis but failed to rescue fertility. It indicates that abortion of oogenesis is not because of mis-regulation of cell death signal but there is oogenesis defect which activates Caspase signaling pathway. Then, we observed *Dyro* mutant and looking for defects which may trigger cell death of germ cells in *Dyro* mutant. We found oogenesis abortion timing is similar to yolk protein mutant but different from amino acid starvation. It suggests that nutrient signal defect does not triggers cell death in *Dyro* mutant. We carefully observed the defect of *Dyro* mutant ovaries and found abnormal morphology of nucleolus and chromosome in nurse cells. It seems chromosome in *Dyro* mutant is thick and nucleolus is limited in small space between thick chromosomes in *Dyro* mutant nurse cells. Other defect we found is aggregated protein accumulation in germ cells. These data suggest that *Dyro* has important role in mid-oogenesis stage germ cell and loss of Dyro causes defect in nuclear of nurse cells which may leads to abortion of oogenesis.

## Introduction

Animals need large amounts of nutrients to make eggs. It is important to make next generation but oogenesis is the big burden for mothers. It is known that *Drosophila melanogaster* has two check points of oogenesis at around stage 2 and 8 (McCall, K. 2004, Pritchett et al., 2009). To avoid consuming nutrients for abnormal eggs which cannot develop, programmed cell death stops oogenesis at check point. It is also known that when female flies are starved, abort oogenesis at mid-oogenesis check point for prioritize mother’s survival. When oogenesis is aborted, caspase cascade is activated, and germ cells of the egg chamber dies (McCall, K. 2004, Pritchett et al., 2009).

Egg chamber of *Drosophila* consists of 15 nurse cells and one egg cell. One of the characteristics of nurse cell is that nurse cells enter endocycle and become polyploid cell. The chromosome structure of nurse cell is dynamically changed during oogenesis (Dej & Spradling, 1999) and mutants which have defects in chromosome disperse are female sterile (King et al, 1981, Keyes and Spradling, 1997). The other significant characteristics of nurse cell is that nurse cell has large nucleolus and transcribe a lot of rRNA (Dapples and King, 1970). During oogenesis, a lot of proteins are synthesized, transported and stored in egg cell which support embryonic development.

It is known that quality control of protein is important for egg. Normally, misfolded proteins are removed by proteasome system or re-folded by chaperon (Hipp et al., 2019, Kandel et al., 2024). Accumulation of aggregated protein can be a stress for cell and be able to induce cell death (Yamada et al., 2023). So aggregated proteins should be removed appropriately.

We found *Dyro* mutant is female sterile and abort oogenesis at around stage8 of oogenesis (Kano and Yagi, 2019). We observed ovaries of *Dyro* mutant using genetic mosaic analysis and we found that loss of *Dyro* in germ line cells is enough to induce abortion of oogenesis. As mid-oogenesis check point exists at around stage 8 (Pritchett et al., 2009), we examined if abortion of oogenesis is regulated by check point mechanism. From observation of cell death timing in the mutant ovary, we found starvation signal is not inducing cell death in *Dyro* mutant, but we found cell death timing is similar to that of the yolk protein mutant. Although timing of oogenesis abortion of *Dyro* mutant is similar, we could not find yolk protein defect in *Dyro* mutant which suggested existence of other developmental defect(s). Thus, we observed *Dyro* mutant ovary further and found morphological defect in nucleolus and chromosome of nurse cell. We also observed protein aggregation in *Dyro* mutant germ line cells, but it is not clear whether protein aggregation is direct reason for oogenesis abortion or one of the results of loss of Dyro function. From these data, we think *Dyro* is involved in the oogenesis regulation and nurse cell nuclear structure might be affected by *Dyro* function.

## Results

### Loss of function mutant phenotype of *Dyro* is female sterile

In our previous study, to show female sterile is loss of function phenotype of *Dyro*, we carried out a jump out experiment of P element using *Dyro*^KG^ allele (Kano and Yagi, 2019). It strongly suggested that female sterility is caused by loss of *Dyro* function by P element insertion but still some uncertainty remains because P element insertion site of *Dyro*^KG^ is intron. To confirm loss of *Dyro* is affecting female sterility, we carried out a series of genetic rescue experiments.

At first, we used two transgenic lines (GRF and FPTB) which have genomic DNA fragment include *Dyro* locus to rescue female sterile phenotype of *Dyro* mutant and both constructs rescued sterility (Table 1). *Dyro*^Δ6^ deletes both *Dyro* and CR45404 (Kano and Yagi, 2019), but genomic rescue construct of GRF does not include CR45404 sequence which indicates that *Dyro* coding genomic region can rescue female sterile phenotype.

**Table 1.** Rescue experiment of female sterile phenotype of *Dyro* mutant.

|  | Larvae+ | egg+, Larvae- | egg- | Total |
| --- | --- | --- | --- | --- |
| GRF/GRF;Δ6/Δ6 | 93 | 10 | 19 | 122 |
| GRF/CyO;Δ6/Δ6 | 92 | 5 | 5 | 102 |
| GRF/GRF;Δ6/TM6B | 102 | 2 | 1 | 105 |
| GRF/CyO; Δ6/TM6B | 94 | 1 | 9 | 104 |
| FPTB/CyO; Δ6/Δ6 | 61 | 0 | 5 | 66 |
| FPTB/FPTB; Δ6/Δ6 | 43 | 3 | 17 | 63 |
| Dyro-Gal4/+; Δ6/TM6B | 107 | 0 | 0 | 107 |
| Dyro-Gal4/+; Δ6/ Δ6 | 0 | 1 | 108 | 109 |
| UASp-Dyro::GFP/ +; Δ6/TM6B | 99 | 0 | 1 | 100 |
| UASp-Dyro::GFP/+; Δ6/Δ6 | 0 | 0 | 105 | 105 |
| Dyro-Gal4/UASp-Dyro::GFP; Δ6/TM6B | 104 | 2 | 3 | 109 |
| Dyro-Gal4/UASp-Dyro::GFP; Δ6/Δ6 | 57 | 16 | 21 | 94 |
| *Δ6/Δ6 | 0 | 0 | 105 | 105 |
| GRF: genomic rescue fragment transgenic |  |  |  |  |
| FPTB: CG6175-GFP.FPTB (BL#83405) |  |  |  |  |
| Δ6: Dyro<sup>Δ6</sup> |  |  |  |  |
| *Kano & Yagi (2019) |  |  |  |  |

**Table 2.** Ovaliole which have egg chamber with more than 15 nurse cells.

|  | More than 15 Nurse cell | Observed ovaliole |
| --- | --- | --- |
| yw | 0 | 502 |
| Dyro<sup>Δ6</sup> | 0 | 617 |
| Dcp1<sup>Prev</sup>; Dyro<sup>Δ6</sup> | 11 | 238 |

Next, we tried to rescue female sterile phenotype using Gal4/UAS system. As female sterile phenotype was rescued by *Dyro* expression by Gal4/UAS system, we conclude that Dyro protein expression is needed for oogenesis (Table 1).

As FPTB and Dyro-Gal4 driven Dyro::GFP expression can rescue female sterile phenotype of *Dyro* mutant (Table 1), tagged Dyro should be expressed in the organ which affects female sterile phenotype. We observed GFP expression of FPTB and Dyro-Gal4/UAS-Dyro::GFP flies (Fig. 1). We detected GFP in nurse cells, egg cell and follicle cells in ovary (Fig. 1 A-F), but Dyro is also expressed in multiple organs in female fly (Fig. 1 G-N). We could observe Dyro expression in egg and nurse cells so it is possible that expression in nurse cell and egg cell is important, but other possibility remains because multiple organs expressing *Dyro*.

**Figure 1.**
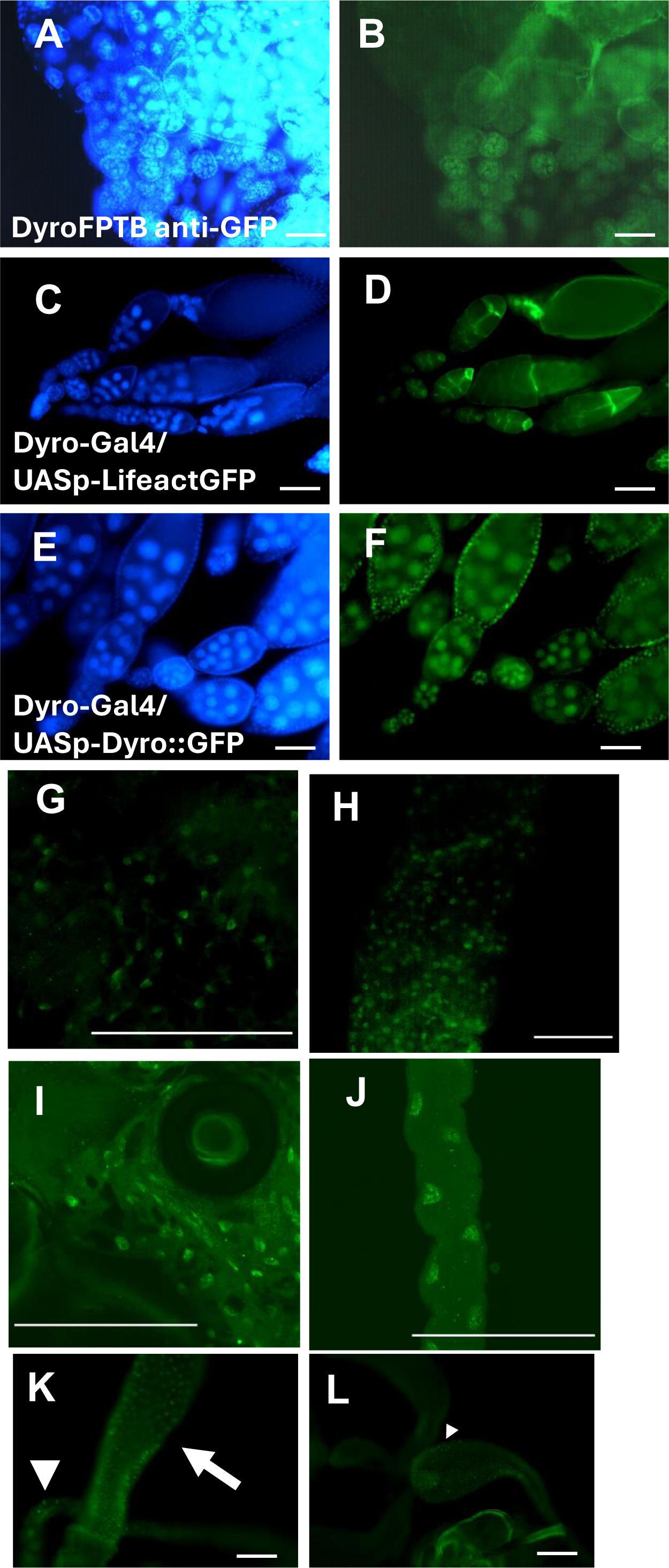
Dyro is expressed in multiple organs in adult fly. To analyze Dyro expression patterns, we used Dyro-GFP.FPTB and Dyro-Gal4, both of which can rescue female sterile phenotype. Ovary (A-F) and some examples of expression in other tissues (G-L). To detect Dyro-GFP.FPTB, we used anti-GFP antibody (B, G-J). For Dyro-Gal4 induced GFP expression, GFP fluorescence was directly observed (UASp-LifeactGFP: D, UASp-Dyro::GFP: F, K,L). GFP was detected in (G) fat body, (H) mid gut, (I) seminal vesicle, (J) Malpighian tubule, (K)mid gut (arrow) and Malpighian tubule (arrowhead), (L) GFP signal was observed in anterior ejaculatory duct (arrowhead). DAPI (Blue), GFP (Green). Bar=100μm.

### Inducing *Dyro* mutant clone egg chambers using FLP-FRT system

We tried germ line mosaic analysis to test *Dyro* expression in germ line cell is important for female sterile phenotype (Fig.2). As marker for recombination was nuclear localized GFP, we could not identify dying egg chamber have *Dyro* mutant cell or not, because nuclear of dying cell is collapsed. But we could observe increase of dying egg chambers in mosaic induced female flies (Fig. 2E). We could also observe debris of dead egg chambers accumulated near the oviduct in mosaic induced ovaries (Fig. 2D). It shows *Dyro* function in germ line cell is important for oogenesis and loss of *Dyro* in germ cell causes abortion of oogenesis.

**Figure 2.**
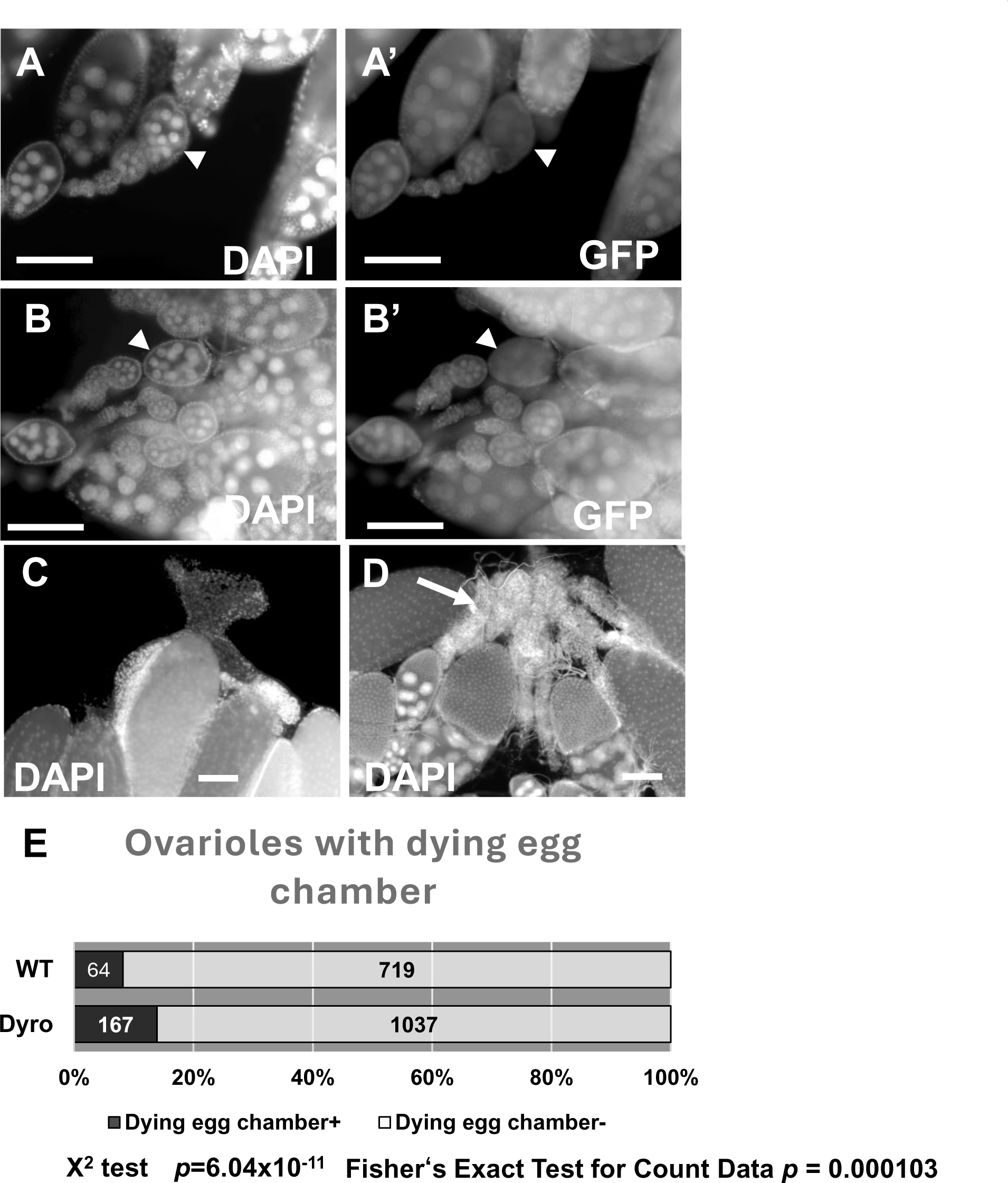
Germ line mosaic analysis of *Dyro*. Germ line mosaic was induced using nanos-Gal4 and UAS-FLP. Ovaries of UASp-FLP; nos-Gal4/+; FRT80B/ubi-GFP FRT80B (A, A’, C) and UASp-FLP; nos-Gal4/+; Dyro FRT80B/ubi-GFP FRT80B (B, B’, D) was observed. Mutant egg chamber which lost GFP expression was observed (arrowhead). In Dyro mosaic ovary, oviduct was filled with dead egg debris (D arrow). (E) Ovarioles with dying/dead egg chamber was counted. Increase of dead egg chamber was observed. Result of the significance test using the Χ-squared test and the Fisher’s Exact Test for Count Data is shown. DAPI (A, B, C, D), GFP (A’, B’). Bar=100μm.

### Cell death in *Dyro* mutant ovary is controlled by Caspase

It is known that Dcp1 is working as effector caspase in the mid-oogenesis check point (Pritchett et al., 2009). If check point to detect developmental defect is killing cells of *Dyro* mutant egg chambers, Caspase cascades include Dcp1 should be working. We examined the effect of *Dcp1* on oogenesis defect of *Dyro* mutant (Fig.3 A-H). In *Dcp1-Dyro* double mutant, strong DAPI signal from fragmented and aggregated DNA of dying cell was not observed (Fig.3 F). But survived cell cannot complete oogenesis and collapsed egg chamber released nuclear of nurse cells into ovariole (Fig.3G). Additionally, some egg chambers have more than 15 nurse cells in *Dcp1-Dyro* double mutant ovary (Fig.3H, Table2). It indicated cell death in *Dyro* mutant ovary is controlled by *Dcp1*.

**Figure 3.**
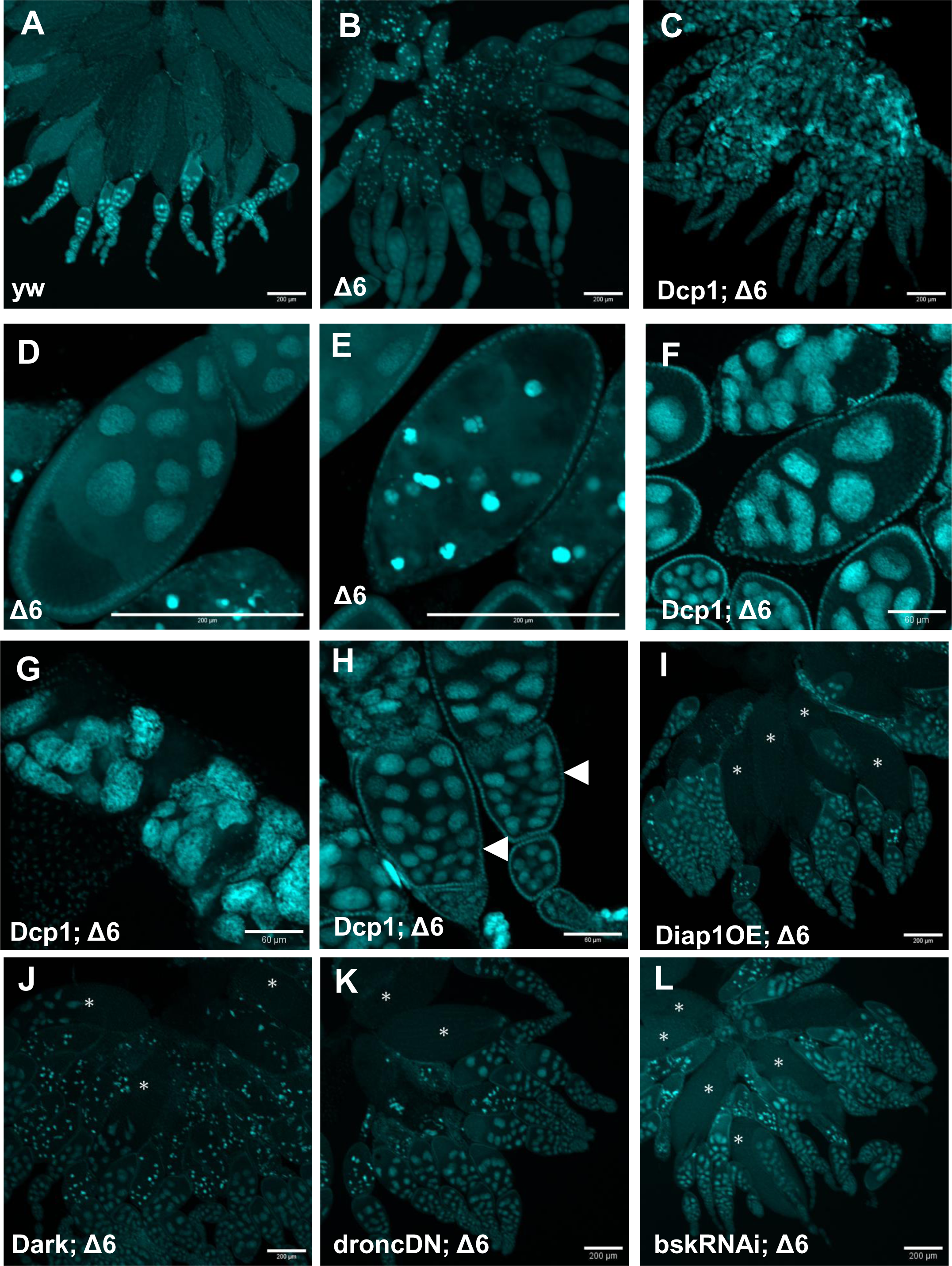
Caspase cascade regulates cell death in ovary of *Dyro* mutant. DAPI stained ovaries were observed. A: yw. B, D, E: Dyro^Δ6^ mutant (Dyro^Δ6^/Dyro^Δ6^). C, F, G, H: Dcp1-Dyro double mutant (Dcp1^prev^/Dcp1^prev^; Dyro^Δ6^/Dyro^Δ6^). I: nos-Gal4/UASp-Diap1; Dyro^Δ6^/Dyro^Δ6^. J: Dark^CD4^/Dark^CD4^; Dyro^Δ6^/Dyro^Δ6^. K: nos-Gal4/UASp-droncCARD; Dyro^Δ6^/Dyro^Δ6^. L: nos-Gal4/bskRNAi; Dyro^Δ6^/Dyro^Δ6^. (I-L) Asterisk shows egg chambers later than stage9. (A) Fertile ovaries with normally developing egg chambers. (B) *Dyro* mutant ovaries have dying egg chambers with strong DAPI signal by fragmented and aggregated DNA. (C) *Dcp1-Dyro* double mutant lose strong aggregated DNA signal but does not have matured egg. (D*) Dyro* mutant egg chamber of stage9 before cell death and (E) dying egg chamber with aggregated DNA. (F) Example of *Dcp1-Dyro* double mutant egg chamber. (G) In *Dcp1-Dyro* double mutant ovary, nuclear of nurse cells leaked from egg chamber to ovariole and accumulated. (H) *Dcp1-Dyro* double mutant have egg chambers with more than 15 nurse cells (arrowhead). Abortion of oogenesis in *Dyro* mutant was suppressed by suppressing cell death by *Diap1* overexpression (I), *Dark* mutation (J), dominant negative *dronc* overexpression (K) and *bsk* RNAi (L). *Asterisk in J-L: egg chambers later than stage9. Bar: 200μm (A-E, I-L), 60μm (F-H)

We genetically manipulated upstream genes of *Dcp1* in *Dyro* mutant background and found oogenesis abortion was suppressed when Caspase activation was inhibited (Table 3, Fig. 3 I-L). It suggests cell death observed in *Dyro* mutant is induced by Caspase pathway activation. As bsk RNAi inhibited cell death in *Dyro* mutant, JNK is involved in Caspase activation in germ line cells. We could rescue cell death in *Dyro* ovary by stopping Caspase pathway activation, but we could not rescue female sterile phenotype. These data suggest *Dyro* mutant has developmental defect which trigger mid-oogenesis check point and even if *Dyro* mutant egg chamber escape cell death, they cannot develop fertile eggs. It is strange that when we suppressed upstream of *Dcp1*, we could observe stage 14 eggs, but we could not observe late-stage eggs in *Dcp1-Dyro* double mutant (Fig.3 C, I-L). It seems some phenotypes are only observed in *Dcp1-Dyro* double mutant. It might be because of genetic background difference or other gene(s) regulated by Caspase signal might be involved.

**Table 3.**
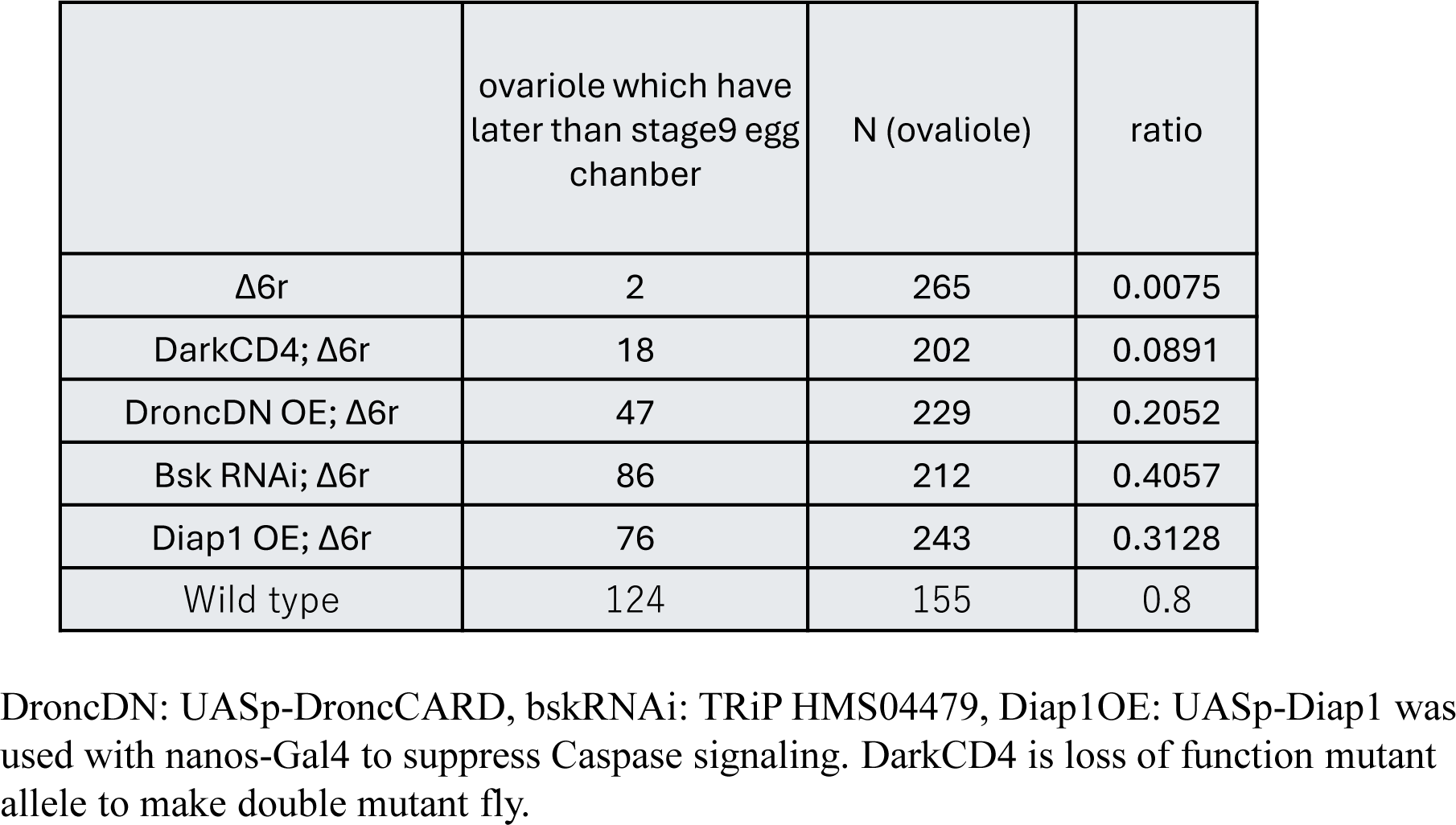
Rescue of cell death in *Dyro* mutant ovary by suppressing cell death signal.

|  | ovariole which have<br>later than stage9 egg<br>chanber | N (ovariole) | ratio |
| --- | --- | --- | --- |
| $\Delta 6r$ | 2 | 265 | 0.0075 |
| DarkCD4; $\Delta 6r$ | 18 | 202 | 0.0891 |
| DroncDN OE; $\Delta 6r$ | 47 | 229 | 0.2052 |
| Bsk RNAi; $\Delta 6r$ | 86 | 212 | 0.4057 |
| Diap1 OE; $\Delta 6r$ | 76 | 243 | 0.3128 |
| Wild type | 124 | 155 | 0.8 |

DroncDN: UASp-DroncCARD, bskRNAi: TRiP HMS04479, Diap1OE: UASp-Diap1 was used with nanos-Gal4 to suppress Caspase signaling. DarkCD4 is loss of function mutant allele to make double mutant fly.

### Abortion of Oogenesis was not related to starvation or yolk protein accumulation

As starvation trigger mid-oogenesis check point, we observed ovary of control and mutant fly fed normal fly food or starved condition (sugar only food). We scored developmental stage of most developed egg chamber in each ovariole and found ovarioles in *Dyro* mutant abort oogenesis at stage 9 but starved fly abort oogenesis at stage 7 (Table 4). It suggests cell death of *Dyro* mutant ovary is not induced by miss fired starvation signal.

**Table 4.** Analysis of abortion timing of oogenesis.

|  | 1 | 2 | 3 | 4 | 5 | 6 | 7 | 8 | 9 | 10 | 11 | 12 | 13 | 14 | total |
| --- | --- | --- | --- | --- | --- | --- | --- | --- | --- | --- | --- | --- | --- | --- | --- |
| Wild type |  |  |  |  |  | 3 | 12 | 6 | 10 | 2 |  | 1 |  | 121 | 155 |
| Δ6r/Δ6r |  |  |  |  |  | 3 | 12 | 41 | 105 | 1 | 5 | 7 | 3 | 5 | 182 |
| Wild type starved |  |  |  |  | 7 | 94 | 50 | 1 | 2 |  |  | 2 |  |  | 156 |
| Δ6r/Δ6r starved |  |  |  |  | 2 | 5 | 144 | 23 | 36 |  |  |  |  |  | 210 |
| C52/Yp1<sup>ts</sup>(29°C) |  |  |  |  |  | 5 | 40 | 36 | 66 | 20 | 1 | 2 | 2 | 42 | 214 |
| C52/Yp1<sup>ts</sup>(25°C) |  |  |  |  |  |  | 2 | 5 | 32 | 34 | 4 | 4 | 5 | 15 | 101 |

The stage of most developed egg chamber in each ovariole from 10 ovaries was recorded except for C52/Yp1<sup>ts</sup>(25°C) (5 ovaries.).)

We searched other defects which could induce abortion of oogenesis. As oogenesis stopped just after yolk accumulation starts, we examined yolk protein mutant phenotype and found yolk protein mutant aborted oogenesis at stage 9 to 10 (Table 4). As oogenesis abortion timing of yolk protein mutant is close to *Dyro* mutant, we examined yolk protein related defect in *Dyro* mutant ovary.

At first, we observed transcription of yolk proteins (Yp1, Yp2, Yp3) in *Dyro* mutant and we could not find difference in transcription of yolk proteins (Fig.4 A-C). Next, we examined yolk proteins in fly hemolymph. We could detect yolk proteins in both control and *Dyro* mutant fly. The amount of yolk protein in *Dyro* mutant hemolymph was higher than control. It might be because mutant ovaries abort oogenesis just after yolk accumulation starts, yolk proteins were not used for oogenesis and accumulated in hemolymph (Fig. 4D). We also observed *yl* expression which encodes receptor for yolk protein uptake and endocytosis activity of egg cell. *yl* gene expression was not reduced in *Dyro* mutant (Fig. 4E). We observed endocytosis activity of oocyte using FM4-64 dye but we could not find endocytosis defect in mutant ovary (Fig. 4F-I). We observed accumulation of GFP tagged Yp1 in Dyro mutant and we could detect Yp1-GFP in *Dyro* mutant oocyte (Fig.4J-L). We carried out a series of yolk protein related experiments, but we could not detect yolk protein defect which led to oogenesis abortion in *Dyro* mutant, so we conclude there is no yolk protein related defects in *Dyro* mutant.

**Figure 4.**
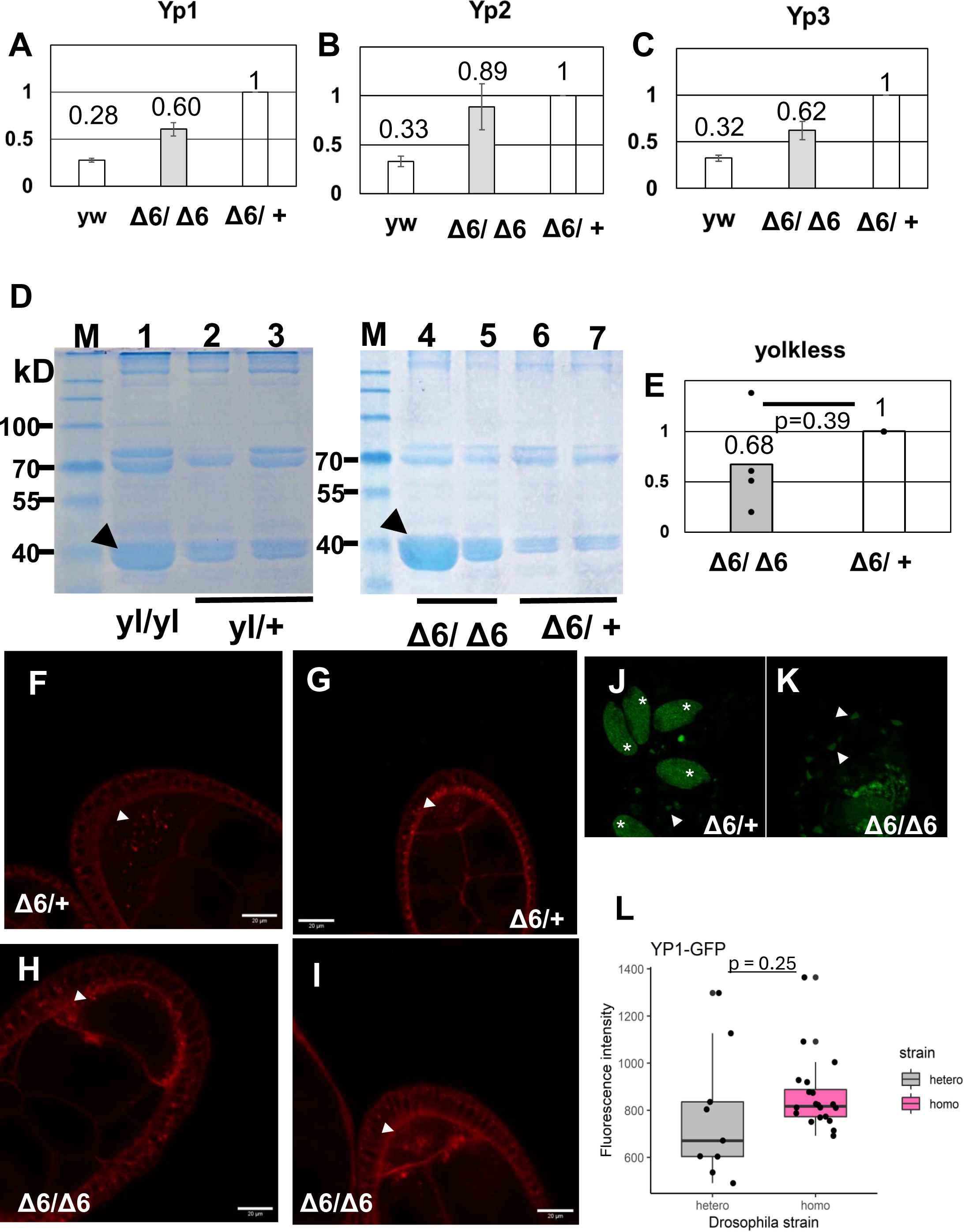
Significant defect was not observed in yolk proteins. (A-C) qPCR analysis of yolk proteins, Yp1, Yp2 and Yp3. Relative quantification between Dyro^Δ6^/+, Dyro^Δ6^/Dyro^Δ6^ and yw. Yolk protein mRNA amount in Dyro^Δ6^/+ was calculated as 1. Comparing hetero and homozygous mutant, yolk protein transcript is reduced in homo but higher than fertile yw strain which indicates that it is within the difference between different strains. (D) Hemolymph of female fly was collected and analyzed by SDS-PAGE and CBB staining. M: molecular weight marker. 1-7: hemolymph sample from indicated genotype female adult flies. Arrowhead shows yolk protein band. For yl/+, Dyro^Δ6^/Dyro^Δ6^ and Dyro^Δ6^/+ 2 independent sample is shown. Yolk proteins were detected in *Dyro* mutant which means yolk proteins are normally secreted into hemolymph. (E)qPCR analysis of *yolkless* expression in *Dyro* mutant. yolkless expression was not affected by Dyro. Relative quantification. n.s.: not significant FM4-64 staining of Dyro hetero fly (F, G) and Dyro mutant (H, I). Fluorescent signal was observed in oocyte (arrowhead). Bar: 20μm. Yp1-GFP was observed in hetero (J) and homozygous (K) Dyro mutant. * eggs of late oogenesis have Yp1-GFP. Arrowheads shows around stage9 oocyte accumulating Yp1-GFP. Fluorescence intensity was measured for stage9 egg cell (L). (E, L) Brunner-Munzel test was used as the significance test.

### Abnormal nucleolus/chromosome morphology was observed in *Dyro* mutant

To find the developmental defect which may cause abortion of oogenesis, we observed *Dyro* mutant ovary and found abnormal nucleolus morphology (Fig.5 A-I). During oogenesis, dynamic change of chromosome morphology is observed. 5-blob was observed in stage 4, and chromosomes are dispersed in later stage and Nuclear of nurse cell at stage 8-9 has large nucleolus (Fig.5 A-C). Although chromosomes of *Dyro* mutant seemed to disperse from 5-blob, but still forming very thick chromosome which looks like polytene chromosome and chromosomes filling nuclear space (Fig.5 D-I). In *Dyro* mutant, we cannot find large nucleolus in stage 9 nurse cell nuclear which is filled with thick chromosomes (Fig.5F, I).

**Figure 5.**
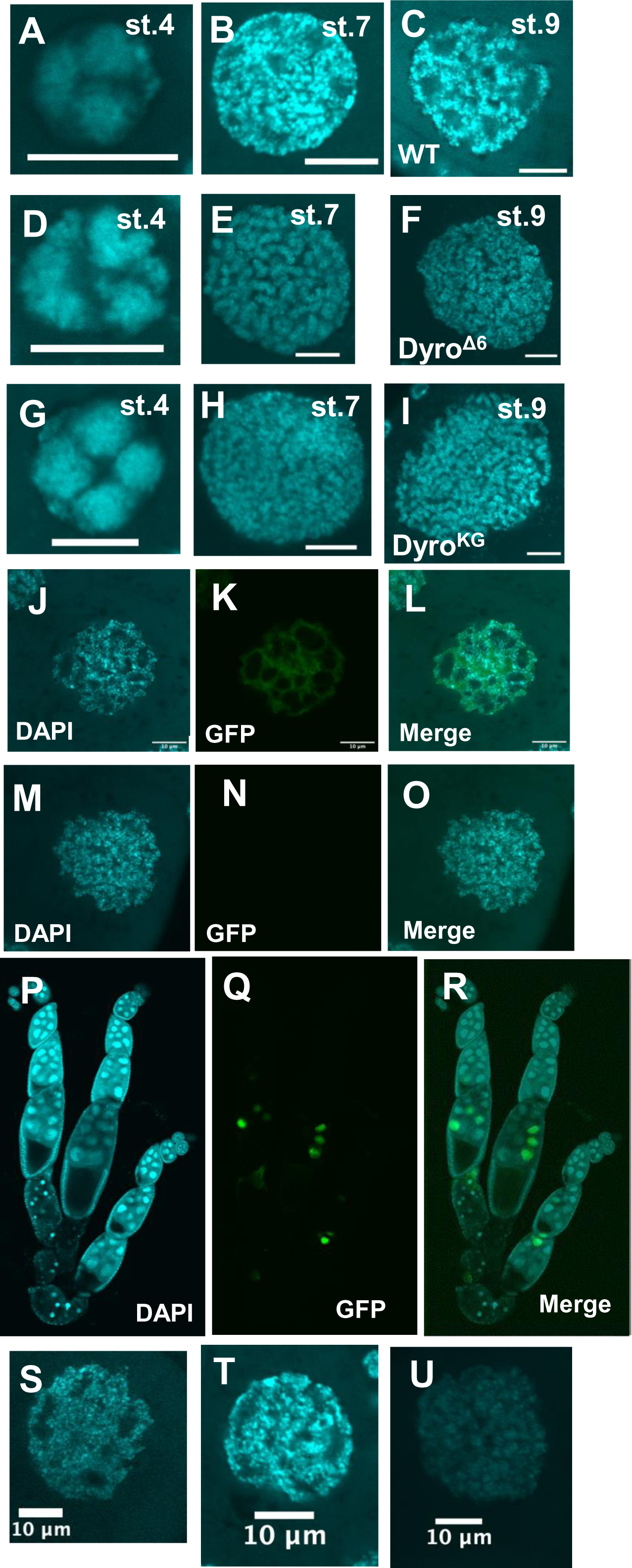
*Dyro* mutant have nucleolus morphology defect in nurse cell. Nuclear of nurse cells are observed. (A-C) Wild type, (D-F) Dyro^Δ6^ mutant, (G-H) Dyro^KG^ mutant. Stage4 (A, D, G), stage 7 (B, E, H), stage9 (C, F, I) nurse cell nuclei are shown. Nurse cell nuclei (J-O) and ovarioles (P-R) of matαGal4/UASp-Dyro::GFP; Dyro/Dyro fly is shown. Only part of nurse cells in single egg chamber expressed Dyro::GFP. Dyro::GFP expressing nurse cell have large nucleoli but nurse cells without Dyro::GFP expression showed mutant like nuclei. Nurse cell nuclei was observed with Dyro knock down flies. (S) CyO/DyroRNAi; Dyro^Δ6^/+ (T) matα-Gal4/DyroRNAi; TM6B/+ (U)matα-Gal4/DyroRNAi; Dyro^Δ6^/Dyro^Δ6^. Nurse cell nuclei phenotype was observed in Dyro knock down with one copy of Dyro (U) but we could not detect the phenotype in Dyro knock down fly with two copy of Dyro (T). Bar: 10μm.

When we used matα-Gal4 to express *Dyro*-GFP, which is known mosaic expression of Gal4 in nurse cells (Rodríguez-Muõz et al., 2022, Fig.5 P-R), rescue of nucleolus/chromosome morphological phenotype was observed only in nucleus which express Dyro-GFP (Fig.5 J-R). The nucleolus defect was also observed in flies with Dyro heterozygous female with Dyro RNAi (Fig.5 S-U). These data suggested that loss of Dyro causes abnormal nucleolus/chromosome morphology in developing nurse cell.

We observed mutant nucleolus using RNA binding fluorescent dye and anti-Fibrillarin antibody (Fig. 6) and found nucleolus exists between small space between chromosomes in *Dyro* mutant. Though it is not clear observed nucleolus is enough to support oogenesis, nucleolus is still existing in *Dyro* mutant.

**Figure 6.**
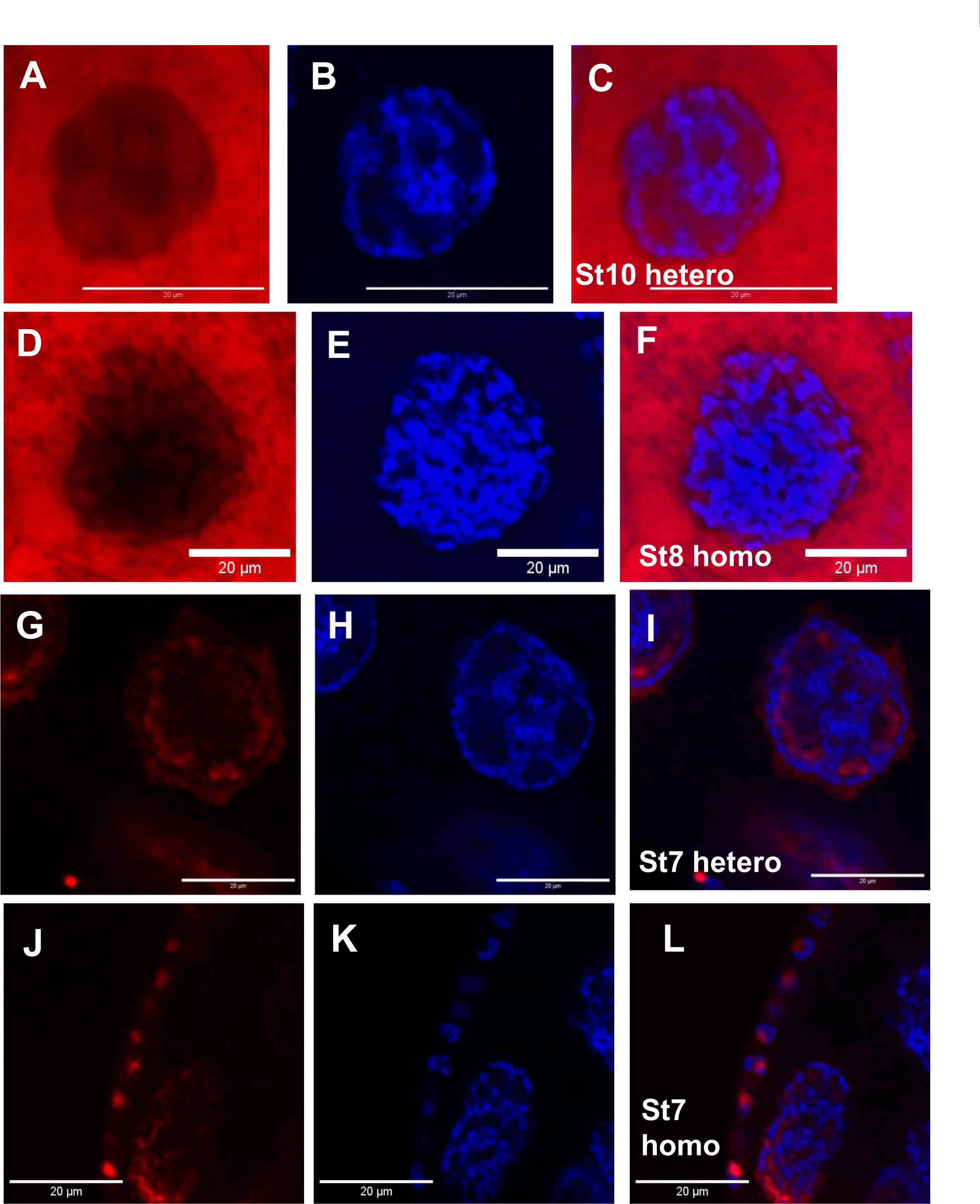
Nucleolus exists small space between chromosomes in *Dyro* mutant nurse cell. Nurse cell nuclear is shown. A-F: Nucleolus Bright Red staining. G-L: anti-Fibrillarin antibody staining. A, D: Nucleolus Bright Red, G, J: anti-Fibrillarin, B, E, H, K: DAPI, C, F, I, L: Merged image. A-C, G-I: Dyro^Δ6^/+. D-F, J-L: Dyro^Δ6^/Dyro^Δ6^. A-C: stage10, D-F: stage8, G-L: stage7 nurse cell nuclear. As nurse cell cytosol contains high concentration of RNA, Nucleolus Bright Red signal is high in nurse cells. Inside nuclear, Nuclear Bright Red signal and anti-Fibrillarin signal was observed between DAPI signal. Bar: 20μm

### Chromosome structure might be affected by Dyro

During oogenesis, endoreplication occurs in nurse cell and chromosome structure is changing after stage6 (Dej & Spradling, 1999). In *Dyro* mutant, it looks chromosome dispersed from 5-blob structure but still have thick polytene like chromosome (Fig. 5). Next, we observed chromosome structure related defect in *Dyro* mutant.

When we observed localization of Dyro in nurse cell, Dyro was observed between nucleolus and DNA. Dyro localization of specific region of chromosome was not observed but does not seem uniform localization (Fig. 7 A-D, G). We examined if Dyro is accumulated in heterochromatin area to interact with heterochromatin structure using anti-HP1 antibody, but we could not find correlation between HP1 and Dyro localization in nurse cell (Fig.7 E-H).

**Figure 7.**
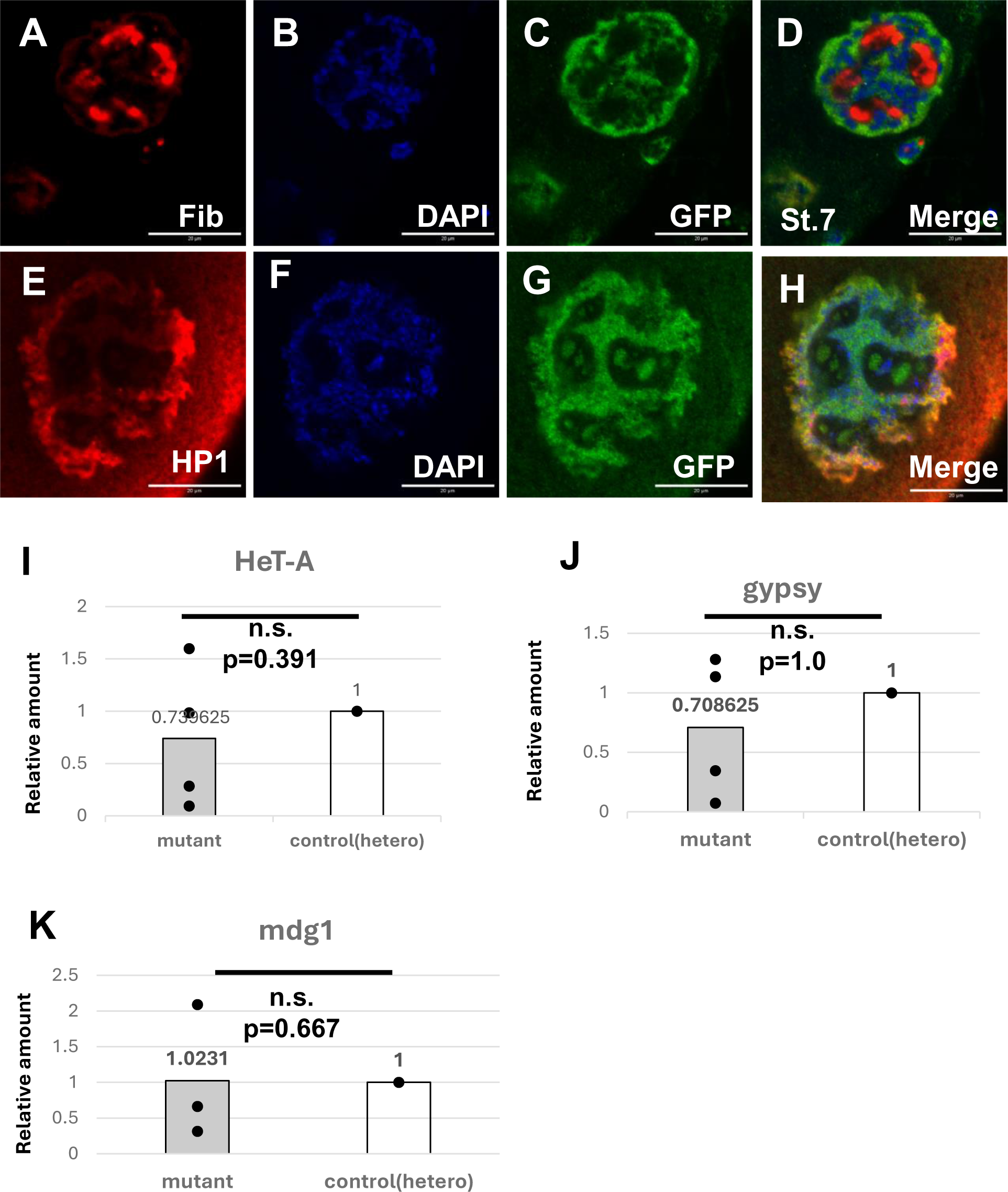
Localization analysis of Dyro in nurse cell nuclear and transposon activity measurement. • A-C: anti-Fibrillarin staining of Dyro-GFP.FPTB. A nuclear of nurse cell is shown. E-H: anti-HP1 antibody staining of Dyro-GFP.FPTB. A nuclear of nurse cell is shown. I-J: qPCR analysis of transposon expression level. I: Expression level of HeT-A. J: Expression level of gypsy. K: Expression level of mdg1. Showing 3 sample data. Data of one sample is excluded as outlier. The data is 43.37 times higher than control. Statistically significant difference was not detected even when including the data. Bar: 20μm. Brunner-Munzel test was used as the significance test.

The other possibility we examined was transposon activity. It is known that de-repression of transposon in germ cell occurs when there are chromosome structure defects and induces oogenesis defect (Sienski et al., 2012, Teo et al., 2018, Moon et al., 2018). To verify possibility that chromosomal defect induces activation of transposon, we monitored transposon activity in *Dyro* mutant. Three transposons were measured but we could not find significant increase of transposon activity (Fig. 7 I-K).

### Aggregated actin was observed in *Dyro* mutant nurse cell

The other abnormality we could detect in *Dyro* mutant ovary was abnormal phalloidin staining signal in germ line cells which suggests existence of aggregated actin in cytosol (Fig.8A, B). Protein aggregation is known as a sign of proteostasis defect. As Proteostasis defect can induce cell death via caspase activation, we tried Proteostat staining of *Dyro* mutant. Slightly stronger signal was observed around nuclear of *Dyro* mutant (Fig. 8C-H) and increased Proteostat signal in *Dyro* mutant was detected (Fig.8I). It is not clear that protein aggregation is directly linked with oogenesis abortion or just indicating unhealthy state of cell, but some protein turnover defect may exist in *Dyro* mutant.

**Figure 8.**
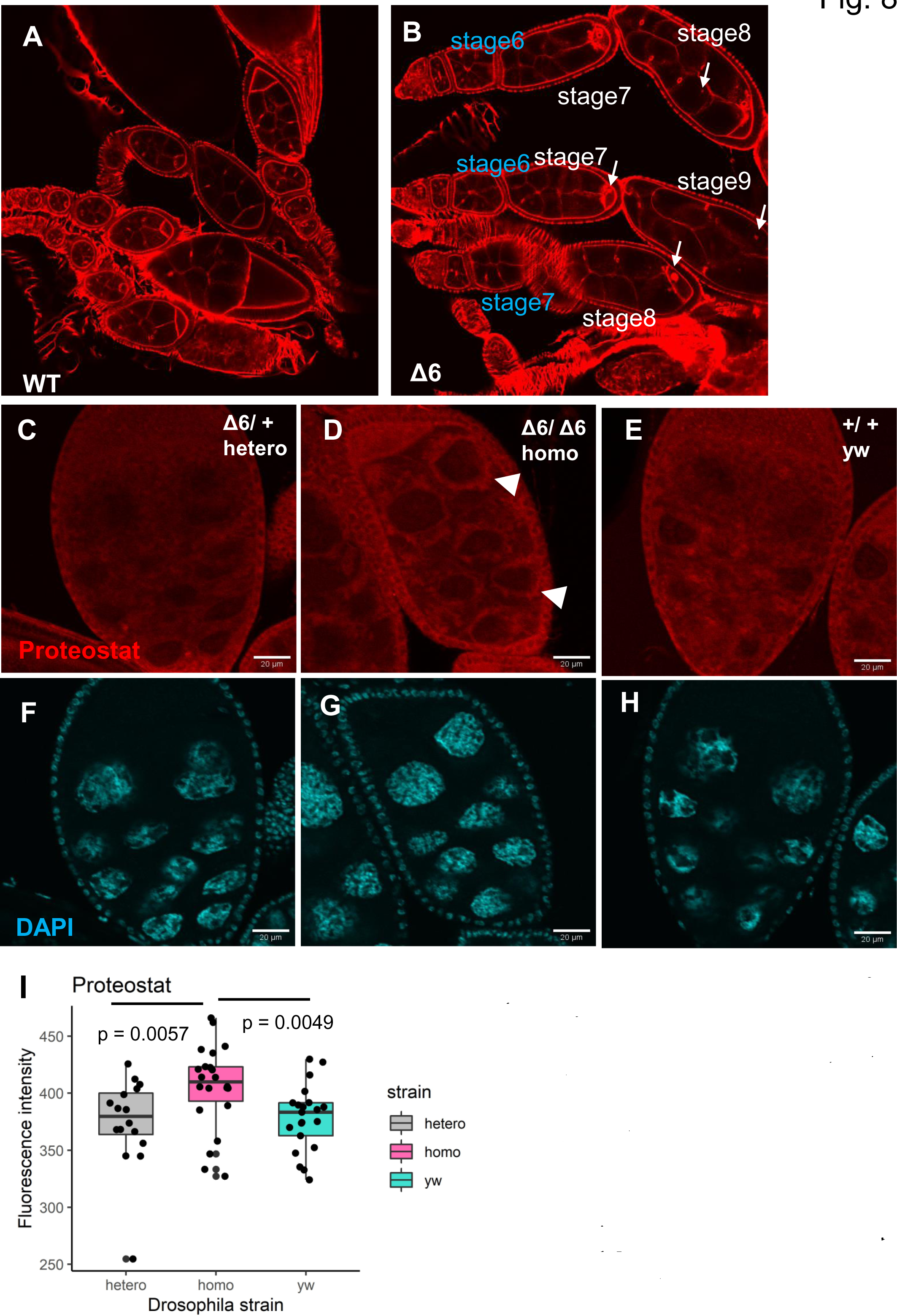
Protein aggregation in germ line cells of *Dyro* mutant. Phalloidin staining of wild type (A) and *Dyro* mutant. In mutant ovary, signals which is not observed in wild type is frequently observed after stage 7 (arrows). Proteostat staining of heterozygous *Dyro* mutant (C), homozygous *Dyro* mutant (D) and control (D) egg chamber. F-G: DAPI signal. (I) Fluorescent intensity of cytosol was measured and analyzed. Brunner-Munzel test was used as the significance test.

## Discussion

Our results show *Dyro* mutant have developmental defect in oogenesis. But the link between observed morphological defects in ovary and female sterile phenotype is still unclear.

From our analysis, it is suggested that abortion of oogenesis is not because of mis-regulated mid-oogenesis check point in *Dyro* mutant. From observation of starved individuals, yolk protein mutant and *Dyro* mutant, there is difference of cell death stage between starvation induced cell death and yolk protein/Dyro mutant. It shows there is difference between starvation induced mid-oogenesis check point and developmental defect induced mid-oogenesis check point. *Dcp1-Dyro* Double mutant analysis shows that mis-regulation of cell death signal is not the reason for abortion of oogenesis, which suggests there is developmental defect which trigger the mid-oogenesis check point in *Dyro* mutant.

One of the possible reasons to abort oogenesis is nucleolus defect because nucleolus is important for producing large amount of rRNA and mRNA for oocyte which support embryonic development. But it is also possible that nucleolus morphological defect was because of abnormal chromosome development in nurse cells and nucleolus existing space between thick chromosomes is enough to support oogenesis.

About 5% of Dcp1-Dyro double mutant ovariole had egg chamber with more than 15 nurse cells. The number was variable and more than 16 (18-35 was observed), it is not mis-differentiation of oocyte like *seh-1* and *mio* (Senger et al., 2011). As the egg chambers with more than 15 nurse cell was not observed in early-stage egg chambers and oocyte was not seen in such egg chamber, it might be defects of cell cycle regulation during oogenesis. It is also possible that dying egg chambers are fused and form abnormal egg chamber. But it is strange that suppressing cell death cause such defect, so it might be because of genetic background.

We observed aggregated protein in *Dyro* mutant nurse cell and oocyte (Fig. 8B). It is a sign that cells have stress, but we could not find it directly trigger cell death or just showing unhealthy state of cells. But it shows *Dyro* mutant have some problem at least stage7.

There are reports that some mutants which have chromosome dispersal defect (Klusza, et al., 2013, Dej and Spradling, 1999, Keyes and Spradling, 1997, King et al., 1981, Koch and King, 1964). In *Dyro* mutant, nurse cell nuclei showed chromosome dispersal from 5-blob but still have thicker polytene chromosome like structure comparing with control and chromosomes occupy most of the space in nuclear. It seems *Dyro* mutant nurse cell nuclear defect is different from known chromosome dispersal defect.

Although significant phenotype was not observed except female sterile, *Dyro* was expressed multiple organs other than ovary (Fig.1 G-N), and when *Dyro* is overexpressed in larval fat body, antimicrobial peptide expression was repressed (Kano and Yagi, 2018). It suggests *Dyro* have unknown minor function in other tissues which is difficult to detect.

## Materials and Methods

### Fly strains

Df(1)C52 (#952), Yp1^ts1^ (#4568), Dyro^KG05833^ (Dyro^KG^, #15102), dcp-1^prev^ (#63814), matα4-Gal-VP16^V2H^ (#7062), nanos-Gal4 (#107748), CG6175-GFP.FPTB (Dyro-GFP.FPTB) (#83405), UASp-FLP (#29730), bskRNAi (#57035, TRiP.HMS04479), DarkCD4 (#23286), UASp-droncCARD (#58992), UASp-Diap1 (#63820) and UASp-LifeactGFP (#58718) was obtained from Bloomington stock center. Ubi-GFP FRT80B (#108348), FRT80B (#106618) was obtained from Kyoto stock center. DyroRNAi (TRiP. HMJ24060) was obtained from NIG-FLY stock center Yp1-GFP (FlyFos019934 [pRedFlp-Hgr] [Yp1[35321]::2XTY1-SGFP-V5-preTEV-BLRP-3XFLAG] dFRT) was obtained from VDRC stock center (#318746). To make transgenic fly using φC31 system, we obtained fly strains from Kyoto stock center (#130431, #130449).

*Dyro^Δ6^*was described in Kano and Yagi (2018)

### Transgenic constructs

#### primers

6175G5A: 5’-AACAACAGATCTGCGGATAGAAAGTATACGGTCG-3’

6175G3A: 5’-CAATTTGCGTCATCGGTACCTTTTGC-3’

GW: 5’-CACCATGTCCAGCAGCGCAGCCACA-3’

GWR: 5’-TTGCATATTGTACAACTGCGGTG-3’

vec-P: 5’-GAATTGGGAATTCGTTAACAGGCCGCTCTAGCCCCCCTCGAATG-3’

vecN6175: 5’-GTGGCTGCGCTGCTGGACATGGTGTGTCGATCAATGAACAGGACCTAAC-3’

6175cVec: 5’-CCGCAGTTGTACAATATGCAATACAAAGTGGTGAGCTCCGCCACCATGG-3’

T-vec: 5’-ATGTGGATCTCTAGAGGTACCTGCAGCCAATCCGCCGCAC-3’

5-fwd: 5’-GAATTCGTTAACAGATCTGCGGCCGCTCTAGAACTAGTG-3’

5-rev: 5’-GTAGCTTCATTGTTGTTGTTGTTGGTAATGG-3’

Gal4_fwd: 5’-AACAACAACAATGAAGCTACTGTCTTCTATC-3’

Gal4_rev: 5’-TTTGCTAGTTTTACTCTTTTTTTGGGTTTG-3’

3-fwd: 5’-AAAAGAGTAAAACTAGCAAATAAACGGG-3’

3-rev: 5’-TAGTGGATCTCTAGAGGTACCTTTTGCATTAATGATTAAAATCC-3’

#### genomic rescue

Genomic region of Dyro was amplified using 6175G5A and 6175G3A. PCR fragment was cut with KpnI and cloned into KpnI-EcoRV cut pBluescriptII SK^-^. Then plasmid with genomic fragment was cut with NotI and KpnI subcloned into NotI-KpnI digested pattB vector.

#### UASp-Dyro::GFP

We made UASp-Dyro::GFP for Gal4 induced expression of *Dyro* in germ cell.

UASp promoter was amplified by PCR with primers vec-P and vecN6175 using pPWH (DGRC) as template.

Dyro cDNA was amplified by PCR with primers GW and GWR using pUAST-CG6175HA (Kano and Yagi, 2019) as template.

GFP + K10 Terminator fragment was amplified by PCR using primers 6175cVec and T-vec using pPWG (DGRC) as template.

PCR fragments and KpnI-BglII cut pattB vector was mixed and assembled using NEBuilder Hi Fi DNA Assembly Master Mix (NEB).

#### Dyro-Gal4

Dyro-Gal4 was generated by removing protein coding region from Dyro genomic rescue construct and insert Gal4 coding sequence.

We employed two strategies to make Dyro-Gal4.

5’UTR and upstream sequence were amplified by 5-fwd and 5-rev primer using genomic rescue construct as template. 3’UTR and downstream sequence were amplified by 3-fwd and 3-rev primer using genomic rescue construct as template. GFP coding sequence was amplified by Gal4_fwd and Gal4_rev primer using pAct5C-Gal4 as template. 3PCR fragment and KpnI-NotI digested pattB plasmid was mixed and assembled by NEBuilder HiFi DNA Assembly Master Mix (NEB). From this reaction, we got a plasmid which have a mutation to change an amino acid of Gal4.

Using genomic rescue construct as template and 3-fwd and 5-rev as template, DNA fragment was amplified by PCR. GFP coding sequence was amplified as described above. Two DNA fragment was assembled by NEBuilder HiFi DNA Assembly Master Mix (NEB). From this reaction, we got a plasmid which have multiple mutations in 5’UTR and upstream region.

We fixed a mutation of first plasmid using DNA fragment from the second plasmid to make Dyro-Gal4 construct. We digested both plasmid by CpoI and XhoI and 2kb fragment from second plasmid was ligated to14kb fragment from first plasmid to fix mutation in Gal4 sequence.

Genomic rescue and UASp-Dyro::GFP construct was introduced into the VK00037 landing site and Dyro-Gal4 construct was inserted into the ZH-51D landing site.

#### Antibodies

Rabbit anti-GFP poly clonal antibody (GenScript A01388) Dilution 1:500 anti-Fibrillarin(38F3) abcam ab4566 Dilution 1:250 (nucleolus marker)

anti-Heterochromatin Protein 1 (HP1) (DSHB, C1A9-s) Dilution 1:50

anti-Rabbit IgG Alexa fluor 488 (Molecular Probe) Dilution 1:500

anti-mouse IgG Alexa fluor 488 (Molecular Probe) Dilution 1:200

anti-mouse IgG Alexa fluor 647 (Molecular Probe) Dilution 1:200

#### Tissue staining reagents

Nucleolus Bright Red (Dojindo laboratories cat.no. N512) was used for nucleolus staining. FM4-64 (Invitrogen Cat. No. F34653) was used for endocytosis assay.

Proteostat (PROTEOSTAT Aggresome detection kit, Enzo Life Science, ENZ-51035-0025) Rhodamine-Phallodin (Molecular Probes R415)

#### Tissue staining

##### Fixation

Adulf female flies were dissected in PBS and ovaries were fixed with 4% formaldehyde/PBS solution for 20minutes. Then samples were washed 5min with PT (PBS with 0.3%Triton X-100) 2 or 3 times and proceed to staining.

##### Staining

For DAPI staining, incubated with PT with DAPI solution for 30minuites, room temperature, protected from light. After staining, wash with PT 5minuites twice and mount with MOWIOL mountant.

For antibody staining, incubate with PT with 0.5% BSA 1hour room temperature as blocking, and incubate with 1st antibody 4°C overnight (2days for anti-HP1). After washing with PT (5min. x2), incubate with 2nd antibody (4°C overnight, protected from light). Then, wash with PT (5min x2) and stain with DAPI. Before observation, sample was mounted with MOWIOL mountant.

For Nucleolus Bright Red staining, incubated with DAPI and 1/1000 diluted Nucleolus Bright Red solution for 30minuites, room temperature, protected from light. After staining, wash with PT 5minuites twice and mount with MOWIOL mountant.

Proteostat staining was carried out following manufacturer’s manual. After fixation, samples were incubated with permeabilizing solution (0.5% TritonX-100, 3mM EDTA, 1x Assay Buffer) for 30minutes. After wash with PT (5min x2), incubate with Proteostat solution (Proteostat reagent 1μL, DAPI stock solution 0.2μL, 1x Assay buffer 500μL) for 30minutes at room temperature, protected from light. Then wash with PT (5min x2) samples were mounted with MOWIOL mountant.

For phalloidin staining, fixed samples were incubated with 1:1000 diluted Rhodamine-phalloidin with DAPI for 30minutes at room temperature. After washing with PT (5min x5) samples were mounted with MOWIOL mountant.

For FM4-64 staining, flies were dissected in PBS and ovaries were cultured in 35mm dish with 2mL of Schneider’s Drosophila medium (Gibco #21720024). Then add 2μL of 10μM FM4-64 and 4μL of DAPI stock solution and incubate 30 minutes under room temperature, protected from light. After incubation, remove culture media and add 250μL of cooled Schneider’s Medium (4°C) and cultured 15 minutes 4°C. Then ovaries were placed on slide glass and mounted with spacer to avoid cover glass crash egg chamber.

Samples were observed using FV3000 confocal microscope (Olympus) or Axioskope2 plus microscope (Zeiss). Images were processed using ImageJ or paint.net.

#### qPCR

##### primers

yp1qF1: 5’-AAGCCCAATGGTGACAAGAC-3’

yp1qR1: 5’-TGTTCGTTGCTCTGATCCTG-3’

yp2qF1: 5’-CATGGGTAATCCCCAATCTG-3’

yp2qR1: 5’-GGTGTAATCGGGCTTGAAGA-3’

yp3qF1: 5’-ACCCTGACCAACTTCAAACG-3’

yp3qR1: 5’-GTTTGGGCGGTGTACTTGTT-3’

ylq1For: 5’-GCACTGCGAGAAGTTCGACAA-3’

ylq1Rev: 5’-TCGAGTCCACGCACTTTCCA-3’

rp49For: *5’*-GACGCTTCAAGGGACAGTATCTG-3*’*

rp49rev: *5’*-AAACGCGGTTCTGCATGAG-3*’*

HeT-AFor: 5*’*-CGCAAAGACATCTGGAGGACTACC-3*’*

HeT-ARev: 5*’*-TGCCGACCTGCTTGGTATTG-3*’*

mdg1For: 5*’*-AACAGAAACGCCAGCAACAGC-3*’*

mdg1Rev: 5*’*-CGTTCCCATGTCCGTTGTGAT-3*’*

gypsyFor: 5*’*-CTTCACGTTCTGCGAGCGGTCT-3*’*

gypsyRev: 5*’*-CGCTCGAAGGTTACCAGGTAGGTTC-3*’*

RNA for qPCR was extracted and purified by RNeasy plus Mini kit (QIAGEN). qPCR reaction was carried out by ABI7500 or StepOne/StepOnePlus Real Time PCR System (ABI) using THUNDERBIRD SYBR qPCR mix (TOYOBO QPS-201) and ReverTra Ace qPCR RT Kit (TOYOBO FSQ-101). RpL32 (rp49) was used as control for each experiment.

##### Hemolymph collection and SDS-PAGE analysis

10 Flies 4day after adult molting was used for each experiment. Flies were prickled with tungsten needle and put on column with nylon mesh. Add 10μL PBS and spin with microcentrifuge (10,000g [rpm?] 1min. 4°C). Add 2xSDS sample buffer to flow through and measured protein amount by Bradford method. Adjust sample volume to contain same protein amount for each lane of SDS-PAGE. Gel was stained with Rapid CBB KANTO 3S (Kanto Chemical).

## Acknowledgement

We thank Kyoto stock center and Bloomington stock center for fly strains. We thank Department of Development and Growth group members (Biological Science, School of Science, Nagoya Univ.) for useful discussion. No research grant supported this work, only supported by funding from university budget.

## Author contributions

T.S., Y.K. and Y.Y. designed and performed experiments and analyzed data. Y.Y. compiled the data and wrote the manuscript with the help of T.S. and Y. K.

